# B cell tetherin: a flow-cytometric cell-specific assay for response to Type-I interferon predicts clinical features and flares in SLE

**DOI:** 10.1101/554352

**Authors:** Yasser M. El-Sherbiny, Md.Yuzaiful Md. Yusof, Antonios Psarras, Elizabeth M. A. Hensor, Kumba Z. Kabba, Katherine Dutton, Alaa A.A. Mohamed, Dirk Elewaut, Dennis McGonagle, Reuben Tooze, Gina Doody, Miriam Wittmann, Paul Emery, Edward M. Vital

## Abstract

**Objective:** Type I interferon (IFN-I) responses are broadly associated with autoimmune disease including SLE. Given the cardinal role of autoantibodies in SLE, we investigated whether a B lineage cell-specific IFN assay might correlate with SLE activity.

**Methods:** B cells and PBMCs were stimulated with IFN-I and IFN-II. Gene expression was scrutinised for pathway-related membrane protein expression. A flow-cytometric assay for tetherin (CD317), an IFN-induced protein ubiquitously expressed on leucocytes, was validated *in vitro* then clinically against SLE diagnosis, plasmablast expansion, and BILAG-2004 score in a discovery cohort (156 SLE; 30 RA; 22 healthy controls). A second longitudinal validation cohort of 80 patients was also evaluated for SLE flare prediction.

**Results:** *In vitro*, a close cell-specific and dose-responsive relationship between IFN-I responsive genes and cell surface tetherin in all immune subsets existed. Tetherin expression was selectively responsive to the IFN-I compared to IFN-II and -III. In the discovery cohort memory B-cell tetherin was best associated with diagnosis (SLE/HC: effect size=0.11, p=0.003;SLE/RA: effect size=0.17, p<0.001); plasmablast numbers in rituximab-treated patients (Rho=0.38, p=0.047) and BILAG-2004. Association were equivalent or stronger than interferon score or monocyte tetherin. The validation cohort confirmed this relationship with memory B-cell tetherin predictive of future clinical flares (Hazard Ratio=2.29 (1.01–4.64), p=0.022).

**Conclusion:** Memory B cell surface tetherin, a reflection of cell-specific IFN response in a convenient flow cytometric assay, was associated with SLE diagnosis, disease activity and predicted flares better than other cell subsets or whole blood assays in independent validation cohorts.

## INTRODUCTION

Type I interferons (IFN-I) are a highly pleotropic group of cytokines that link the innate and adaptive immune systems and play a pivotal role in autoimmune disease(1-3). All nucleated cells express IFN-I receptors and will express a set of interferon-stimulated genes (ISGs) after exposure to IFN-I(4, 5). Hundreds of effects of IFN-I on various cellular processes, interactions and disease processes have been described. A challenge in the assessment of IFN-I response in an individual disease is therefore ensuring the appropriate cellular response can be detected within this complex system.

IFN-I proteins are unstable in blood and not easily detected even in monogenic interferonopathies with known high IFN-I production, possibly due to their efficient binding to the abundant IFN receptor(6). IFN-I activity is therefore usually measured using expression of ISGs in whole blood. We previously analysed in ISG expression in sorted cells from SLE (a prototypic IFN-medicated disease) and healthy individuals and reported that ISG expression was markedly higher in monocytes compared to other circulating immune cells, which therefore dominates ISG assays that use unsorted blood(7).

These differing levels of ISG expression in cell populations may be due to the rate of turnover of each population, their trafficking to sites of higher IFN-I production in inflamed tissues, or priming for IFN-I response by other inflammatory mediators. In autoimmunity, IFN-I assays may have value to predict flares, and response to a range of different targeted therapies(8). However, existing whole blood IFN biomarkers show poor or uncertain correlation with disease activity(9-11).

The measurement of IFN-I status using whole blood ISG expression has two key weaknesses in interpreting pathogenic processes. First, changes in expression may reflect expansion or contraction of certain circulating leukocyte populations(12, 13) that differ in their level of ISG expression. This characteristically occurs in inflammatory diseases. In the case of SLE, lymphopenia is almost universally seen(14). So any difference in whole blood gene expression may not necessarily indicate a change in production or exposure to IFN-I.

Second, analysing whole blood ISG expression does not allow detection of key pathogenic processes among the noise of other, less relevant, effects of IFN-I on biology. For example, B cells are a key mediator in SLE(15, 16). IFN-I stimulates B cells to differentiate into plasmablasts which are expanded in SLE and correlate with disease activity(17, 18). We previously demonstrated that the rate of plasmablast regeneration post-rituximab predicts clinical outcome(19). We also previously showed that IFN-I imprints plasma cells for the secretion of the proinflammatory molecule ISG15(17). Assessment of IFN-I activity in unsorted blood, gives limited information about the degree to which B cells have specifically been stimulated by IFN-I. Further, gene expression assays do not prove that a phenotypic change in target cells has occurred – there has been no widely used biomarker for IFN response at a protein level.

In order to resolve these problems, we developed a flow cytometric assay to allow measurement of IFN-I response in individual cells without the need for cell sorting. We measured the expression of Tetherin (*BST2*, CD317), a GPI-anchored protein with unique topology which is ubiquitously expressed on the surface of nucleated cells. This molecule is prominent in viral immunology and encoded by a commonly-measured ISG expressed in all leucocytes (4, 5, 20-22). Unlike most ISGs, *BST2* encodes a cell surface protein and can be easily measured in patient samples by flow cytometry. Siglec-1 is another flow cytometric IFN-I biomarker previously described(23, 24). However, Siglec-1 is only expressed on monocytes so resolves the issue of changes in size of cell populations but does not allow interrogation of individual cell subset IFN-I responses, including the key B lineage cell populations that are strongly linked to clinical and experimental disease.(25-27)

We hypothesised that a dominant pathogenic role of IFN-I in SLE is its effect on B-cells, promoting plasmablast differentiation and clinical disease. Using in vitro stimulation and sorted cells from SLE patients and healthy individuals we showed that Tetherin accurately captures cell-specific differences in response to IFN-I. An crucial issue in biomarker research is demonstrating that biomarkers are predictive, correlate with a range of outcomes and can be reproduced in validation studies. In our studies, longitudinal analysis of discovery and validation cohorts showed that memory B cell tetherin more accurately correlates with plasmablast expansion, clinical features of disease and predicted flares better than monocyte tetherin or whole blood ISG expression.

## MATERIALS AND METHODS

The SLE discovery cohort samples included 156 consecutive SLE patients as well as 22 age-matched healthy controls (HC) and 30 active ACPA-positive ANA-negative RA patients (DAS28) =3.9, 95% (CI 3.23–4.56) as a non-SLE inflammatory disease control. An independent validation cohort consisted of 80 SLE patients recruited and studied longitudinally (total n=236). Disease activity was assessed at the time of sampling using the British Isles Lupus Assessment Group (BILAG-2004)(28). Validation cohort patients were also followed up for subsequent flare (BILAG A or B). SLE patients’ demographics and disease activity are shown in (Table S1). Patients with acute or chronic viral infection at the time of blood sampling were excluded from this study. All individuals provided informed written consent and this research was carried out in compliance with the Declaration of Helsinki. The patients’ blood samples used for this study were collected under ethical approval, REC 10/H1306/88, National Research Ethics Committee Yorkshire and Humber–Leeds East, and healthy control participants’ peripheral blood was collected under the study number 04/Q1206/107. All experiments were performed in accordance with relevant guidelines and regulations. The University of Leeds was contracted with administrative sponsorship.

The full details of methods see online Supplementary file and a previously published methodology paper(7).

## RESULTS

### BST2/Tetherin as a cell-specific phenotypic biomarker of IFN-I response

Global gene expression profiles showed that many ISGs were responsive to both IFN-α and IFN-γ while other ISGs responded specifically to IFN-α (7, 17). We therefore tested the influence of IFN-α (IFN-I) and IFN-γ (IFN-II) on 31 of most common reported ISGs by qPCR analysis (TaqMan) on B-cells *in vitro* as previously described (17) (Supplement figure S1). Results from *in vitro* stimulation confirmed that *BST2* was in the ISG group predominantly responsive to IFN-I.

For this reason, we applied multi-parameter flow cytometry analysis to detect and quantify the tetherin on PBMCs described in supplementary methods. We used a gating strategy allowing to define: T-cells, NK-cells, monocytes as well as B-cell subsets: naïve, memory and plasmablasts. For each of these populations mean fluorescence intensity (MFI) of tetherin is shown compared to isotype control (Figure 1A). We compared cell-surface tetherin protein levels by flow cytometry versus *BST2* gene by qPCR for these six FACS-sorted cell subtypes from ten SLE patients and six HC (total n=16). *BST2* gene expression levels were substantially positively correlated with tetherin protein levels on the cell surface within each of the subtypes (Figure 1B). These data confirm that varying level of tetherin/*BST2* expression between cell subsets and differences between individuals may be captured using flow cytometry without the need for cell sorting. Furthermore, we compared tetherin MFI on monocytes, B-cells and T-cells with Siglec-1 in samples from 25 SLE patients and 5 healthy controls. We confirmed that tetherin correlated with Siglec-1 only on monocytes because other cell subsets lack expression of Siglec-1 (Figure S2).

**Figure 1.**
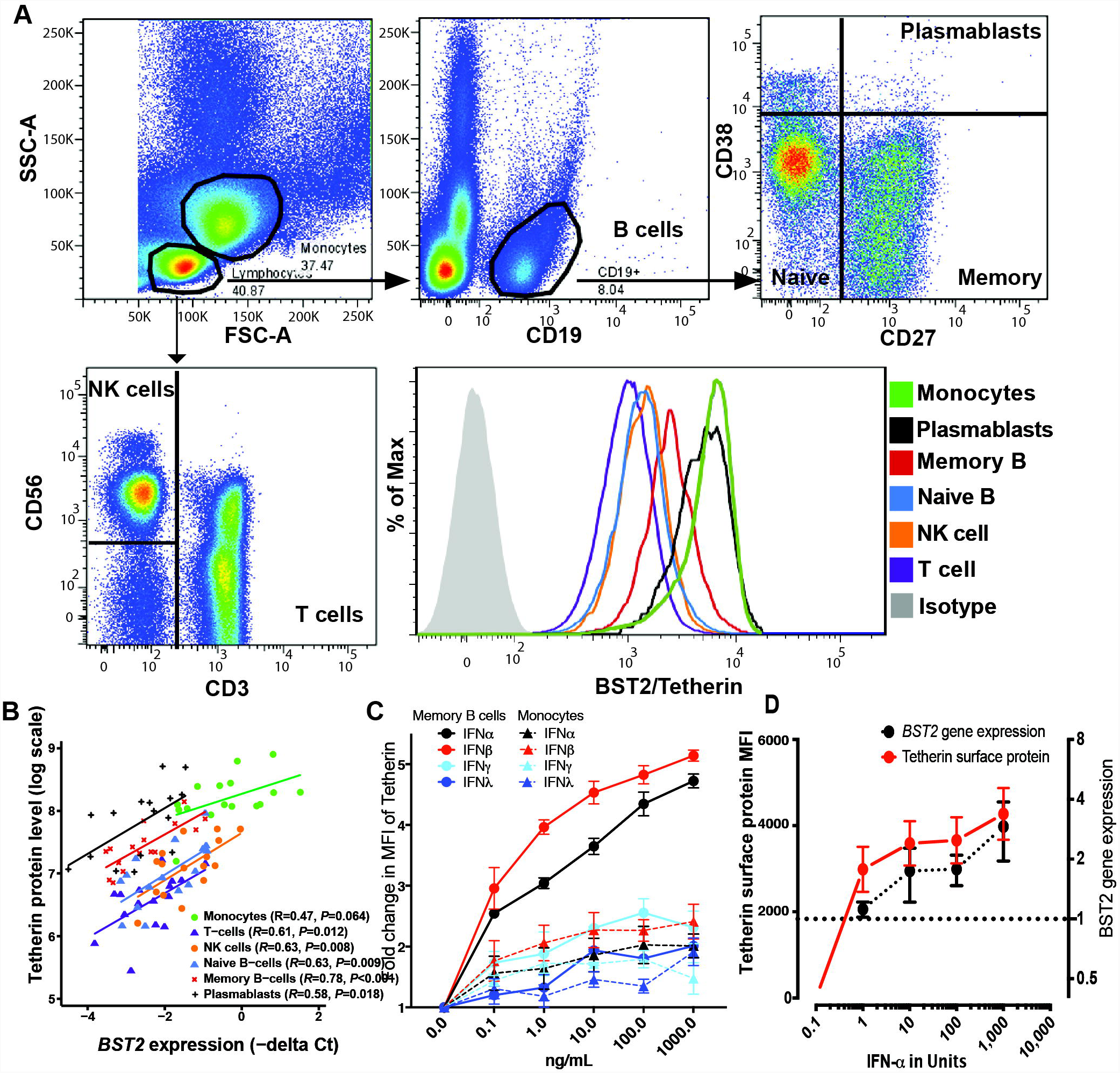
Tetherin is a scalable cell specific measure of IFN-I response. **(A)** Gating strategy for flow cytometric assessment of tetherin on immune cell subsets. An example flow cytometry analysis plot for analysis of tetherin protein expression on individual immune cell subsets is shown. Forward (FSC-A) and side scatter (SSC-A) were used to define lymphocytes and monocytes. B-cells were defined as CD19+ lymphocytes and subdivided into naïve, memory and plasmablast subsets using CD27 and CD38. T-cells were defined as CD3+ and NK-cells as CD3-CD56+ lymphocytes. The mean fluorescence intensity (MFI) of tetherin is shown compared to isotype control. **(B)** In order to validate tetherin as a cell specific marker, tetherin protein expression was compared with expression of its gene *BST2* in various immune cell subsets. Unsorted PBMCs from the samples presented in Fig 1/Table S5 were analysed using flow cytometry for surface tetherin on each subset (y axis) and correlated with gene expression data for *BST2* from the sorted subset (x axis). There was a strong correlation between gene expression and protein within each subset allowing differences in ISG expression between cell subsets to be measured without-cell sorting (Monocytes R=0.47, P=0.064; T-cells R=0.61, P=0.012; NK cells R=0.63, P0.008; Naïve B-cells R=0.63, P0.009; Memory B-cells R=0.78, P0.001; Plasmablasts R=0.58, P0.018) **(C)** HC PBMCs were stimulated with increasing doses of IFN-α, IFN-β, IFN-γ, and IFN-λ then each subset was evaluated with flow cytometry for mean fluorescence intensity of tetherin of memory B-cells (solid circles, solid curves) and Monocytes (Triangles, dashed curves) (n=3) **(D)** Sorted B-cells were stimulated in vitro with increasing doses of IFN-α and then evaluated with flow cytometry for mean fluorescence intensity of tetherin (red line) as well as expression of its gene, *BST2.* We found a parallel increase in each marker

### Dose response of tetherin to IFN-I, IFN-II and IFN-III

We tested dose-response relationship of tetherin on all circulating cell subsets towards IFN-α, IFN-β (both IFN-I), IFN-γ (IFN-II), and IFN-λ (IFN-III). HC PBMCs were stimulated for 48h with doses ranging from 0.1–1000 ng/mL then analysed by flow cytometry. Interestingly, tetherin MFI on memory B-cells was the most responsive to increasing doses of IFN-α and IFN-β, with more modest response to IFN-γ and IFN-λ. Although monocytes had the highest expression of tetherin in patient samples and the highest basal expression in unstimulated HC PBMCs, they showed much lower fold change in tetherin response to IFN-I stimulation (Figure 1C). Furthermore, purified B-cells response curves for *BST2* gene expression and tetherin protein MFI revealed a dose response closely matched response to IFN-α (Figure 1D). We concluded that tetherin MFI by flow cytometric analysis could accurately measure change in expression of *BST2* in response to IFN-I and could be used to capture IFN-I exposure in a dose- and cell-specific manner.

### B-cell surface tetherin protein levels best demonstrate disease-associated IFN response in SLE

We next compared tetherin protein expression in immune cell subsets in SLE patients and HC to determine which cell subset would demonstrate disease-associated change in IFN response best. Using all discovery cohort data from SLE and HC, tetherin protein levels were compared between groups across cell subtypes by flow cytometry results summarised (Table 1, upper panel).

**Table 1:** Tetherin levels in Cell Subsets in SLE patients and controls

| All patients | SLE<br>n=113 | Within-SLE, between | HC<br>n=17 | Between-group | Between-group, between |
| --- | --- | --- | --- | --- | --- |
|  |  | SLE subtypes |  | SLE:HC | subtype |
| | | Ratio (90% CI), $P$ value | | Ratio (90% CI), $P$ value | Ratio (90% CI), $P$ value |
| Monocytes | 3388 | Reference | 2837 | 1.19 (0.91, 1.58), $P = 0.293$ | Reference |
| T-cells | 687 | 0.20 (0.19, 0.22), $P < 0.001$ | 475 | 1.45 (1.17, 1.79), $P = 0.005$ | 1.21 (1.00, 1.46), $P = 0.092$ |
| NK-cells | 1129 | 0.33 (0.31, 0.36), $P < 0.001$ | 824 | 1.37 (1.09, 1.72), $P = 0.024$ | 1.15 (0.94, 1.40), $P = 0.258$ |
| Naïve B-cells | 1118 | 0.33 (0.30, 0.36), $P < 0.001$ | 712 | 1.57 (1.22, 2.03), $P = 0.004$ | 1.32 (1.03, 1.68), $P = 0.062$ |
| Memory B-cells | 1586 | 0.47 (0.43, 0.50), $P < 0.001$ | 1033 | 1.53 (1.23, 1.92), $P = 0.002$ | 1.29 (1.04, 1.58), $P = 0.046$ |
| Plasmablasts | 2650 | 0.78 (0.72, 0.84), $P < 0.001$ | 1813 | 1.46 (1.15, 1.86), $P = 0.009$ | 1.22 (0.99, 1.51), $P = 0.115$ |
| Rituximab<br>naïve only | SLE<br>n=76 | Within-SLE, between | HC<br>n=17 | Between-group | Between-group, between |
|  |  | SLE subtypes |  | SLE:HC | subtype |
| | | Ratio (90% CI), $P$ value | | Ratio (90% CI), $P$ value | Ratio (90% CI), $P$ value |
| Monocytes | 3206 | Reference | 2949 | 1.09 (0.80, 1.48), $P = 0.657$ | Reference |
| T-cells | 666 | 0.21 (0.19, 0.23), $P < 0.001$ | 494 | 1.35 (1.07, 1.70), $P = 0.034$ | 1.24 (1.01, 1.52), $P = 0.080$ |
| NK-cells | 1068 | 0.33 (0.31, 0.36), $P < 0.001$ | 857 | 1.25 (0.98, 1.58), $P = 0.129$ | 1.15 (0.93, 1.41), $P = 0.271$ |
| Naïve B-cells | 1132 | 0.35 (0.32, 0.39), $P < 0.001$ | 740 | 1.53 (1.20, 1.95), $P = 0.004$ | 1.41 (1.15, 1.73), $P = 0.006$ |
| Memory B-cells | 1574 | 0.49 (0.45, 0.53), $P < 0.001$ | 1074 | 1.47 (1.17, 1.83), $P = 0.005$ | 1.35 (1.11, 1.64), $P = 0.013$ |
| Plasmablasts | 2597 | 0.81 (0.74, 0.89), $P < 0.001$ | 1885 | 1.38 (1.08, 1.76), $P = 0.033$ | 1.27 (1.02, 1.57), $P = 0.068$ |

Values under SLE and HC show age-adjusted mean tetherin protein levels compared within SLE patients and between SLE patients and controls. The tetherin cell protein data were ln-transformed prior to analysis, therefore the back-transformed results represent ratios of values in each of the subtypes relative to monocytes within SLE patients, and the ratio of SLE patients to HC in each of the subtypes. Interaction ratios express the ratio of the extent of the difference between SLE and HC in each subset relative to monocytes

Tetherin levels differed significantly between cell subtypes within SLE, being highest for monocytes, 20% of monocyte level in T-cells, and 78% of monocyte level in plasmablasts (all p<0.001). Comparing between SLE and HC groups showed that tetherin MFI on monocytes did not significantly differ from HC (SLE:HC ratio 1.19, P=0.293), whereas a significant higher level was seen in in SLE for all other subsets (ratios 1.37–1.57, all P<0.05). Comparing the between group ratio of each subset against that for monocytes showed that the disease-associated increase of memory B-cell tetherin showed the greatest difference than monocytes’ (P=0.046).

Rituximab treated SLE patients could confound accurate measurement of B-cell phenotype in these patients. We therefore repeated these analyses in rituximab-naïve patients (Table 1, lower panel). In rituximab-naïve patients (n=58), the largest disease-associated increase in tetherin expression was seen in naïve (1.53) and memory B-cells (1.47). These ratios for naïve and memory B-cells were significantly different from monocytes (P=0.006 and P=0.013 respectively). These results indicate that differences in IFN response at a protein level between cell subsets are clinically relevant and B-cell tetherin is the most clinically relevant parameter.

### Clinical validation of Tetherin IFN assay: diagnosis

We compared the performance of the tetherin flow cytometric assay in distinguishing between patients with a diagnosis of SLE, active RA, or HC. Given our previous results, for this analysis we only included rituximab-naïve patients controlled for age (Figure 2). Full statistical table is shown in Supplement (Table S3). We considered effect sizes to be small (0.01), medium (0.06) or large (0.14) as described by Cohen (29). Tetherin data revealed a marked difference between cell subsets. Monocyte tetherin did not differentiate SLE from healthy control at all with ratio 1.19 (0.87– 1.61) and effect size 0.03. T-cells and NK-cells had moderate effect sizes of 0.06 each. However, naïve and memory B-cell subsets had medium to large effect size of 0.11 with ratios of 1.63 (1.26 –2.11) and 1.59 (1.21–2.09) respectively.

**Figure 2.**
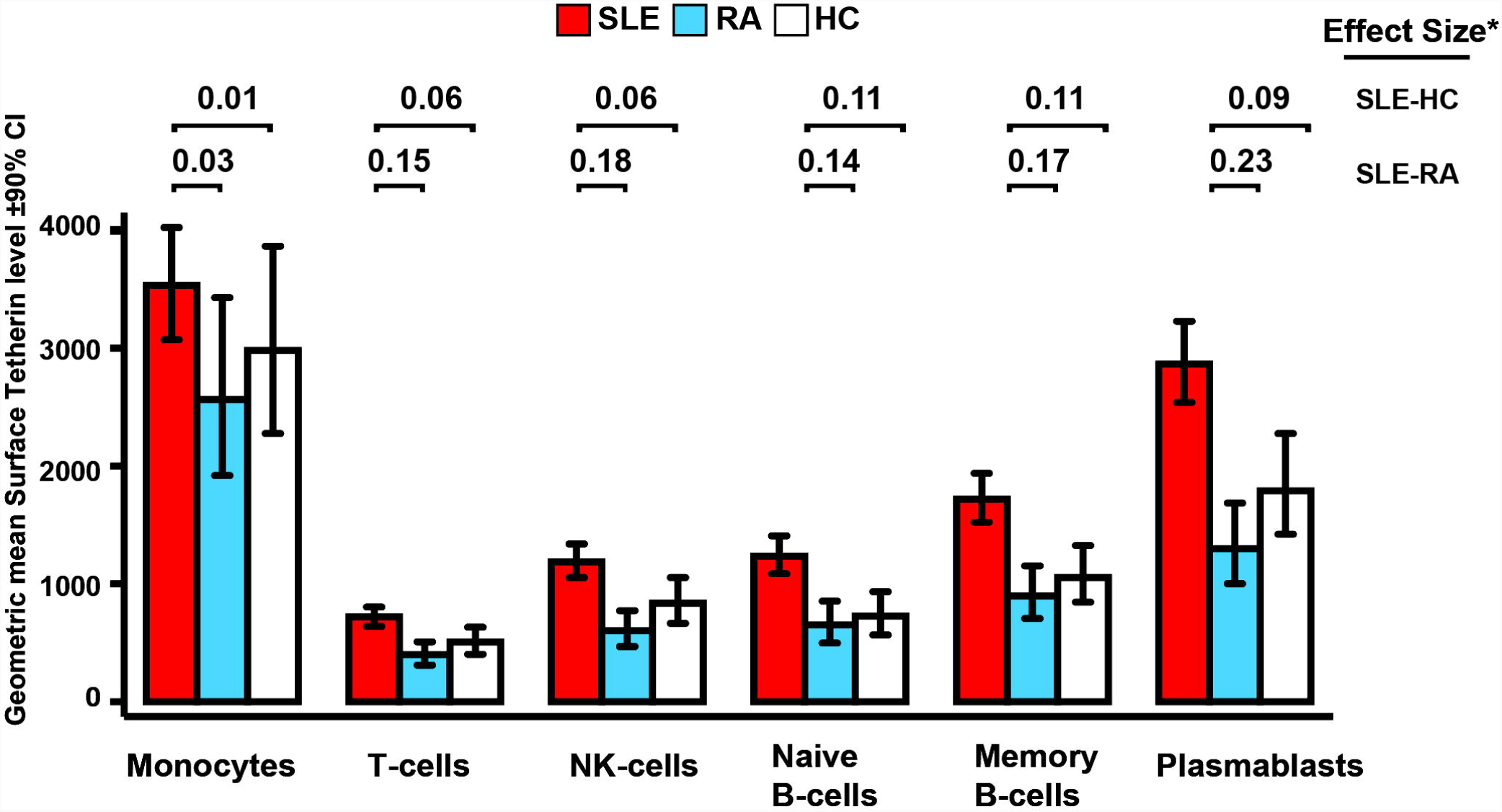
Comparison of tetherin flow cytometric IFN assay against diagnosis. Age-adjusted differences between patients with SLE (red) and patients with active RA (DAS28>3.2; blue) or healthy controls (white). cell surface BST2/tetherin protein levels from cell subtypes identified through flow cytometry of PBMCs. Effect sizes (partial eta squared) indicate which of the variables differed to the greatest extent between the different groups. We considered effect size 0.01 to be small, 0.06 to be medium and 0.14 to be large (29).

Tetherin was able to differentiate SLE from other inflammatory disease when compared with active RA. Tetherin on monocytes had no diagnostic function with ratio 1.37 (1.00–1.88) and effect size 0.03. However, all other cell subsets had moderate to large effect sizes ranging from 0.14 to the largest for plasmablasts at 0.23, with ratio 2.20 (1.66–2.93).

### Clinical validation of IFN assays: disease activity

For disease activity, we investigated the association between the number of active organ systems (BILAG domains scoring A, B or C) per patient compared to tetherin on cell subsets as well as our recently described IFN score A, which comprises of 12 IFN-I selective ISGs (7). We controlled for age in all SLE patients (164 observations in 124 patients). The number of active domains was categorized 0 (n=22), 1 (n=54), 2 (n=57) or ≥3 (n=31).

At the 10% level of significance, disease activity was associated with IFN Score A (R^2^ = 0.08, P=0.027) and tetherin surface expression on T-cells (R^2^=0.07, P=0.007), NK-cells (R^2^=0.09, P=0.001), memory B-cells (R^2^=0.09, P=0.006) and plasmablasts (R^2^=0.06, P=0.020). The degree of association was weaker and hence non-significant for monocytes (R^2^ = 0.04, P=0.179) and naïve B-cells (R^2^=0.04, P=0.103).

For IFN Score A, the relationship with disease activity was not linear. The only significant association between the score and disease activity was attributable to patients with the most severely active disease (≥3 domains). A similar, although non-significant, pattern was observed for tetherin on monocytes. In contrast, there was a linear relationship between memory B-cell tetherin and disease activity with a stepwise increase in expression for each increase in number of active domains (Figure 3A).

**Figure 3.**
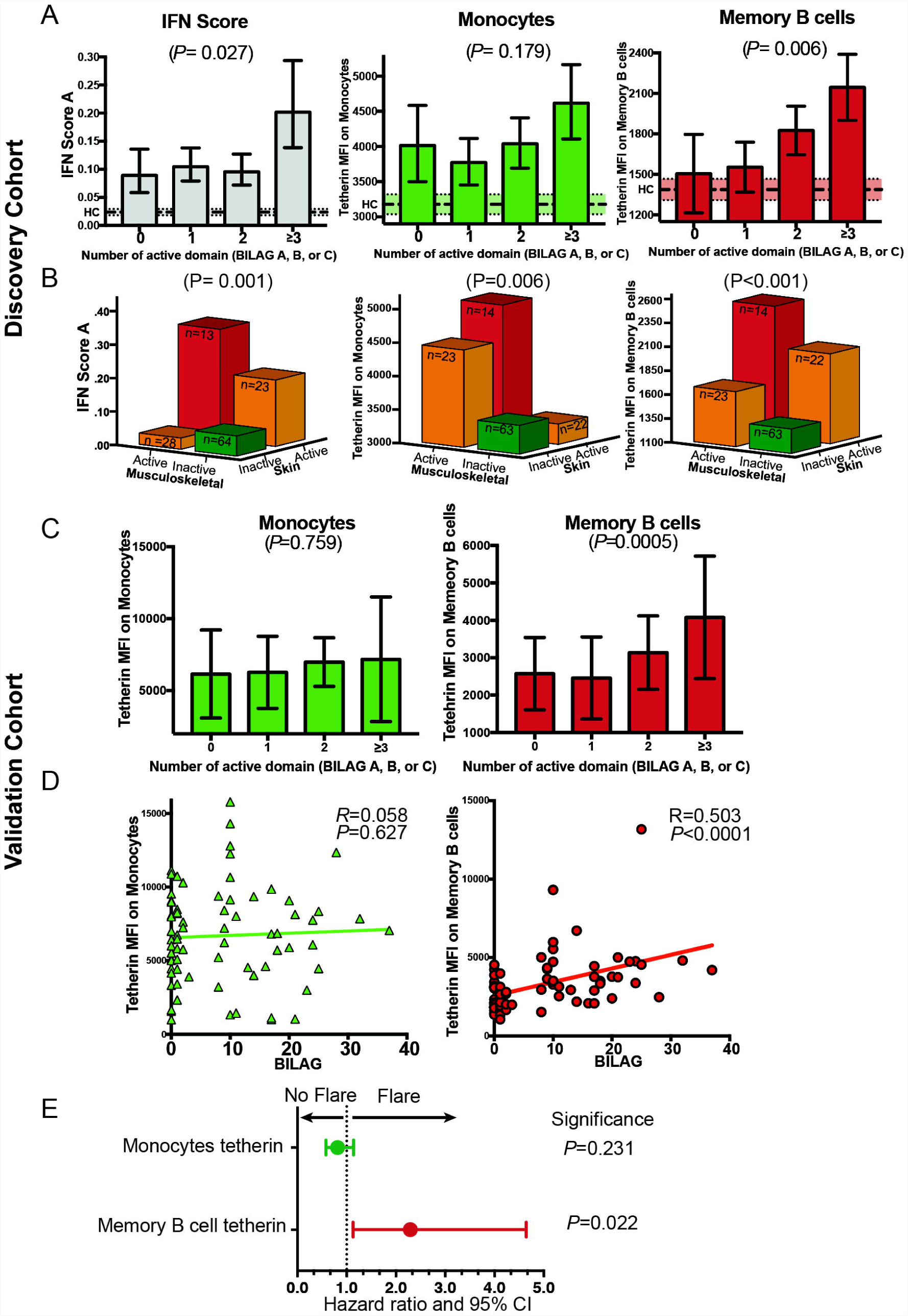
Association between IFN assays and disease activity in SLE. **(A)** Discovery cohort: association between different IFN assays and number of organ systems (domains) with active disease 164 observations in 124 SLE patients). IFN score shown as 2-dCT (i.e. taller bars represent higher expression) with 90% CI. Dotted lines and shaded areas represent mean and 90% CI of 23 healthy controls (HC). Interferon Score was increased in patients with ≥3 active domains but not in patients with 1 or 2 active domains compared to 0 (remission). Tetherin measured on memory B-cells demonstrated a more consistent stepwise increase with increasing disease activity. **(B)** Discovery cohort: association between different IFN assays and musculoskeletal and mucocutaneous disease activity. Disease activity was defined as Active (BILAG A or B) or Inactive (BILAG D or E). Patients with activity in other organs were excluded. For Interferon Score there were inconsistent relationships with disease activity, with an increase with skin involvement, but not musculoskeletal involvement alone. For monocyte tetherin increased protein expression was seen with musculoskeletal disease activity but not for skin activity alone. Tetherin measured on memory B-cells demonstrated a consistent relationship with both common types of clinical disease. **(C)** Validation cohort (n=80): scatter plots show association between overall disease activity (BILAG Global score) and tetherin. There was a significant association between BILAG Global score and tetherin on memory B-cells, but not with tetherin on monocytes. Bar charts show that association between tetherin and number of active organ domains was similar to the discovery cohort, analysed as in (A). **(D)** Tetherin on monocyte tetherin did not (Hazard Ratio=0.814, 95%CI 0.580-1.141, P=0.231) while tetherin on memory B-cells significantly predicts subsequent clinical flare (Hazard Ratio=2.290, 95% CI 1.013–4.644, P=0.022).

To investigate whether the difference between IFN-assays was because of the type of organ system affected, we analysed the two most commonly affected domains (mucocutaneous and musculoskeletal) in combination, excluding patients with activity (BILAG A, B or C) in any of the other domains. Although there was a significant relationship between each IFN-assay and overall disease activity, the relationship with IFN-assays varied between these two organ systems (Figure 3B). For IFN Score A, increased expression was only seen with mucocutaneous disease activity. While, for tetherin on monocytes, increased level was observed only in patients with musculoskeletal disease activity. This may explain why this assay does not show a linear relationship with disease activity. However, memory B-cell tetherin had a more consistent relationship with disease activity in both organ systems. The level of tetherin was lowest in patients in clinical remission, higher in patients with active disease in a single organ, and highest in patients with active disease in both organs.

Numbers of patients with other active organ domains were more limited. Of patients with no activity in other domains, 12 had active (A or B) haematological disease (immune mediated haemolysis or thrombocytopenia). Memory B-cell tetherin MFI on the active and inactive haematology groups was 1954 vs. 1494 respectively, *P*=0.005. Eight patients had active renal disease. Comparing these 8 active renal patients versus inactive disease also revealed a significant increase in tetherin on memory B-cells (tetherin MFI 2625 vs. 1562, P=0.005).

### Clinical validation of IFN assays: plasmablasts

Lastly, in the discovery cohort, we used plasmablast count to represent current B-cell activity and differentiation. IFN-I is known to promote the differentiation of memory B-cells into plasmablasts (30). We have previously shown that post-rituximab early rapid population of plasmablasts led to an early clinical relapse(31, 32). We hypothesized that the memory B-cells tetherin level would correlate with circulating plasmablasts numbers post-rituximab reflecting an increased rate of differentiation secondary to IFN-I. Results are shown in Table 2. In rituximab-naïve patients, no relationship was revealed between any tetherin IFN-assay and plasmablast count, while in post-rituximab patients there was no correlation with tetherin on monocytes, NK or T-cells. But memory B-cell tetherin was significantly correlated with numbers of plasmablasts after (Spearman’s R=0.38, P=0.047) (Figure 4A) as well as inversely correlated with time to clinical relapse (R=0.623, P=0.022) (Figure 4B).

**Table 2:** Association between candidate IFN assays and plasmablast level following B-cell depletion therapy

|  | Plasmablast Count (cells x 10<sup>9</sup>/L) |  |
| --- | --- | --- |
|  | Pre-rituximab<br>(*n* = 50) | Post-rituximab<br>(*n* = 28) |
| Interferon Score A | -0.11, *P* = 0.448 | 0.24, *P* = 0.219 |
| Tetherin protein level: |  |  |
| Monocytes | -0.08, *P* = 0.592 | 0.20, *P* = 0.296 |
| T-cells | -0.16, *P* = 0.269 | 0.32, *P* = 0.096 |
| NK-cells | -0.14, *P* = 0.324 | 0.05, *P* = 0.795 |
| Naïve B-cells | -0.04, *P* = 0.801 | 0.30, *P* = 0.121 |
| Memory B-cells | 0.07, *P* = 0.618 | 0.38, *P* = 0.047 |

Values are Spearman's Rank Correlation Coefficient and *P* values for correlation between plasmablast count and Interferon Scores A and B (measured on unsorted PBMCs) as well as tetherin MFI on each cell subset (analysed using flow cytometry).

### Independent Validation Cohort

The independent validation cohort consisted of a further 80 patients with SLE recruited and studied prospectively. Memory B-cell and monocyte tetherin levels were measured using fresh lysed whole blood in an independent accredited diagnostic laboratory. Disease activity was measured at the time of sampling using BILAG-2004. Patients were followed up for subsequent flare (BILAG A or B).

We found a similar relationship between tetherin and disease activity as in our discovery cohort. For memory B-cell tetherin, there was a significant relationship between number of organ domains and active disease (P=0.0005) but no relationship with monocyte tetherin (P=0.759, Figure 4C). There was a significant association between global BILAG score and memory B-cell tetherin (Spearman’s R=0.503, P<0.0001) but no association with monocyte tetherin (R=0.058, P=0.627). Additionally, in this cohort we demonstrated that in patients in clinical remission at the time of sampling (n=36), memory B-cell tetherin predicted time to clinical flare. In multivariable cox-regression analysis including memory B-cell tetherin, monocyte tetherin and age, memory B-cell tetherin was a significant predictor of subsequent BILAG A/B flare (Hazard Ratio=2.290, 95% CI 1.013–4.644, P=0.022). Monocyte tetherin did not significantly predict flare (Hazard Ratio=0.814, 95%CI 0.580-1.141, P=0.231). In conclusion, we independently confirmed that disease activity is related to IFN-I response in memory B-cells measured using tetherin, and further, that this is predictive of clinical outcome.

## DISCUSSION

In this study we have demonstrated the value of a novel cell-specific biomarker based on the IFN-inducible protein tetherin, using *in vitro* methods and clinical studies in humans. We showed that flow cytometric measurement of cell surface tetherin on memory B-cells captures cell-specific IFN-I response, is dose-responsive, and has a strong and consistent relationship with disease activity, B-cell activity and time to flare in two SLE cohorts. These results are important because IFN-I and B cells have a role in many autoimmune diseases and their measurement has potential to stratify outcomes and use of therapies, but previous studies yielded conflicting results(33).

Better biomarkers are needed in SLE. EULAR Treat-to-target recommendations advise treating to a target of low disease activity, while minimising exposure to glucocorticoids(34). Predictors of a severe disease trajectory or flares are needed to achieve this. Response to conventional and targeted therapies in SLE and related diseases is variable and re-classification of autoimmune diseases according to pathogenic mechanisms instead of clinical features has been proposed(33).

The crucial role for IFN-I in the pathogenesis of SLE and related diseases is indicated by genetic susceptibility and monogenic interferonopathies as well as evidence of over-expression(33). As such, it has face validity as a stratification biomarker. Existing studies indicate the potential value of measuring IFN-I in diagnosis and prediction of flares. IFN-I biomarkers may also predict clinical response to TNF-blockade, B cell depletion and IFN-I-blockade in RA and SLE(33).

Nevertheless, there are limitations to previous approaches to measuring IFN-I activity and some previous results have been contradictory. Direct measurement of IFN-I protein is limited by the number of different ligands and instability in serum, with most cell types expressing the type I IFN receptor. A recent improvement was the use of single-molecule arrays (Simoa). The higher sensitivity of Simoa allows reliable measurement of IFN-α (6). However, this is currently expensive and limited in availability and has not been validated against clinical outcomes. When using ISG expression based methods, another issue is the effect of other IFN subtypes or other inflammatory mediators. ISGs are known to fall into distinct subsets, which may be due to the effect of IFN-II (7, 9). We previously showed that there are different patterns of ISG expression in different autoimmune diseases. In the present paper we confirmed that tetherin is selectively responsive to IFN-I and we included ANA-negative RA as an inflammatory disease control.

While candidate biomarker discoveries in autoimmunity are numerous, a significant challenge is in validation in clinically relevant contexts (35). An important aspect of our work is the degree of pre-clinical and clinical validation. We present validation against longitudinal endpoints. Our data demonstrate good face and construct validity, as well as concurrent and prospective criterion validation and feasibility in a routine clinical setting.

Cell specific measurement based on flow cytometry has been demonstrated previously using expression of Siglec-1, another cell surface protein convenient for flow cytometry that is expressed by monocytes. Monocyte Siglec-1 has been shown to correlate with disease activity as well as predict autoimmune congenital heart-block (24, 36, 37). This was a significant advance in analysis of interferon status. In the present work we advance this principle further by using a marker expressed on all circulating cells. Tetherin captures the same information as Siglec-1 on monocytes, but also evaluates other cell subsets. We have shown that results from these different subsets vary, with the strongest clinical correlation for memory B cells. This has distinct advantages when there is particular interest in a specific cell population, such as with the B cell directed therapies rituximab and belimumab in SLE. B cell response to IFN-I is crucial in SLE.

While there were many associations between tetherin protein expression and clinical features of SLE, tetherin on memory B cells seemed to be particularly important. This marker correlated best with clinical features, and was the only marker to be associated with plasmablast number. After B cell depletion with rituximab, there is a highly variable rate of plasmablast repopulation that predicts clinical relapse. Understanding the determinants of these repopulation patterns may reveal upstream factors controlling B cell autoreactivity. One previous paper reported a relationship between serum BAFF titres and numbers of plasmablasts at relapse(38). However, BAFF may not be the only factor. IFN-I also promotes B cell activation and differentiation into plasmablasts and plasma cells (27, 39). This may include direct influences, for example in animal models IFN-I influences BCR and TLR-mediated response to self-nuclear antigen. Our work provides data in the human disease to support this observation(40, 41). Additionally, IFN-I induces a plasma cell phenotype that secretes ISG15 with additional pro-inflammatory effects(17). In the present study we found that memory B cell tetherin correlated with plasmablast expansion after rituximab. A plasmablast signature was recently shown to be a strong biomarker for SLE and we and others previously showed that plasmablast expansion after rituximab was strongly predictive clinical relapse(19, 42, 43).

The tetherin biomarker has some limitations. First, although this flow cytometric assay avoids confounders that may affect ISG expression scores, analysing a single interferon-inducible transcript may be more susceptible to the influence of other inflammatory stimuli, which we cannot exclude based on these results. However, our data comparing SLE with the RA disease control are very consistent with those we observed using interferon scores with a clear difference in IFN Score A and Tetherin expression between SLE and RA. Tetherin, like all IFN-I biomarkers, may be influenced by acute or chronic viral infections, which was excluded from this study. It may be more difficult to perform flow cytometry in some situations. However, with widespread use of flow cytometry in cell-targeted therapies in autoimmunity and oncology as well as in routine monitoring of HIV, addition of tetherin cell surface staining is a highly cost-effective test. Tetherin may be analysed in combination with B-cell and plasmablast flow cytometry to stratify both B-cell and IFN-I blocking therapy.

In summary, we describe measurement of the interferon-inducible protein tetherin on B cells as a cell-specific biomarker with a number of advantages and widespread applications in clinical and laboratory research in this rapidly expanding area of immunology.

## Acknowledgments

We thank C.F. Taylor for gene expression analysis and A. Droop for database organization

## AUTHOR CONTRIBUTIONS

All authors were involved in drafting the article or revising it critically for important intellectual content, and all authors approved the final version to be published. Dr. Ed Vital had full access to all of the data in the study and takes responsibility for the integrity of the data and the accuracy of the data analysis.

### Study conception and design

El-Sherbiny, Emery, Vital.

### Acquisition of data

El-Sherbiny, Yusof, Psarras, Hensor, Kabba, Mohamed, Tooze, Doody.

### Analysis and interpretation of data

El-Sherbiny, Yusof, Psarras, Hensor, Dutton, Tooze, Doody, Elewaut, McGonagle, Wittmann, Emery, Vital.

## Notes

**Funding:** This work was supported by the National Institute for Health Research (NIHR) Leeds Musculoskeletal Biomedical Research Unit. The views expressed are those of the author(s) and not necessarily those of the NHS, the NIHR or the Department of Health. A.P. is funded by University of Leeds 110 Anniversary Research scholarship. A.A.A.M. was funded by Egyptian government scholarship. E.M.V. is funded by NIHR Clinical Scientist Fellowship (CS-2013-13-032) and the Academy of Medical Sciences (grant AMS-SGCL8).

**Disclosures / conflicts of interst:** The authors declare that they have no competing interests. Data and materials availability: All data are contained within the manuscript and the Supplementary Materials.

