## Supplementary materials for "B cell tetherin: a flow-cytometric cell-specific assay for response to Type-I interferon predicts clinical features and flares in SLE"

**Supplementary Materials and Methods**

### Patients

Ethical approval for the study was given by Leeds East National Research Ethics Service Committee (REC 10/H1306/88). For the discovery cohort, consecutive SLE patients were recruited from a large tertiary referral centre with no selection for disease activity or therapy. All patients were positive for anti-nuclear antibodies using the Bioplex 2000 assay. All SLE patients met ACR/SLICC 2012 criteria. All RA patients met ACR/EULAR 2010 criteria [52]. Disease activity was assessed using BILAG-2004, a validated index that scores SLE-associated disease activity separately for each organ domain [39]. Current and previous medications for SLE were recorded. Absolute lymphocyte count was obtained from a routine clinical laboratory full blood count and used to calculate absolute counts for flow cytometry subsets. The discovery cohort consisted of a further 80 patients recruited to the same ethically approved study. In this cohort flow cytometry was performed in a routine clinical diagnostic laboratory using lysed whole blood with the same antibody clones as the discovery cohort. These patients were assessed using BILAG-2004 at the time of sampling as well as rate of clinical flare (new BILAG A or B) in follow up.

### Samples

Peripheral blood mononuclear cells (PBMCs) were separated using density gradient method (Lymphoprep^TM^, Alere Technologies, Norway) from EDTA anticoagulated peripheral blood, isolated cells were washed twice by Dulbecco's Phosphate-Buffered Saline (DPBS) then labeled with panel of monoclonal antibodies for immunophenotyping or FACS cell sorting.

### Antibodies

The following antibody clones were used in this study; CD19 (clone HIB19); CD69 (clone FN50); CD56 (clone B159), CD3 (clone SK7), CD4 (clone RPA-T4); CD8 (clone SK1); CD27(clone M-T271), all from BD Biosciences (Oxford, UK); CD14 clone TÜK4); CD16 (clone Clone VEP13), CD38 (clone DX9); CD64 (clone 10.1.1), CD169-Siglec-1 (clone 7-239), all from Miltenyi Biotec (Bisley, UK) and BST2/tetherin/CD317 (clone 26F8) from eBiosciences (Hatfield, UK).

### Flow cytometry and cell sorting

Cell surface expression of bone marrow stromal cell antigen 2 (BST2/tetherin/CD317) was determined using flow cytometry analysis of stained peripheral blood mononuclear cells (Figure 3). Flow cytometry was performed for the discovery cohort and experimental stream using a Becton Dickinson (BD) LSRII flow cytometer at flow cytometry facility, Wellcome trust Brenner building, or BD FACSCanto for validation cohort samples at the diagnostic immunology service laboratory service, Leeds Teaching Hospitals NHS Trust. both performed using BD FACSDiva software.

The following antibodies were used for flow cytometry: Tetherin/CD317 (PE-labeled RS38E clone; BioLegend), CD56 (APC-labeled clone REA196; Miltenyi Biotec), CD38 (PE-Vio770-labeled REA572; Miltenyi Biotec), CD19 (VioBlue-labeled LT19; Miltenyi Biotec), CD8 (PerCP-labeled BW135/80; Miltenyi Biotec), CD27 (VioBright-FITC-labeled M-T271; Miltenyi Biotec) CD4 (APC-Vio770-labeled VIT4; Miltenyi Biotec), CD3 (VioGreen-labeled BW264/56; Miltenyi Biotec) and CD14 (FITC TÜK4, Miltenyi Biotec), CD27 (Alexa flour700 M-T271, Becton Dickenson), CD69 (FITC FN50, Becton Dickenson). We used a gating strategy allowing to define viable single cell events of lymphocytes and monocytes. B-cells were defined as CD19+ lymphocytes and subdivided into naïve (CD19+CD27-CD38Dim), memory (CD19+CD27+CD38+) and plasmablast (CD19+CD27++CD38++) subsets, while, T-cells were defined as CD3+ lymphocytes and NK-cells as CD3-CD56+ lymphocytes. For each of these populations mean fluorescence intensity (MFI) of tetherin is shown compared to isotype control. For 25 SLE patient samples and 5 healthy controls we also evaluated tetherin (CD317) and SIiglec-1 (CD169) on CD19+ B cells, CD3+ T cells and CD64+CD14+ monocytes.

For FACS cell sorting a six-way sort was performed on a BD Influx™ cell sorter to individual pure lymphocyte subsets (T-cell CD3+, NK-cells CD3-CD56+, B-cell memory CD19+CD27+CD38dim, naïve B-cells CD19+CD27-CD38dim, plasmablasts CD19+CD27+CD38++, monocytes CD14+). Purity of each sorted population was confirmed before RNA extraction using flow cytometry (>98% purity for each subset) as well as gene expression of corresponding lineage markers.

### Gene expression studies

Total RNA purification kit (Norgen Biotek, Canada) was used to extract RNA either from PBMCs or sorted cell subsets. To ensure the purity of total RNA, genomic DNA was eliminated from all samples using DNAse according to manufacturer’s instructions and quality were tested using RT+/RT– experiment. For cDNA synthesis from total RNA acquired, Fluidigm^®^ Reverse Transcription Master Mix buffer was used according to manufacturer’s instructions and protocols including a mixture of random primers and oligo dT were used for priming.

TaqMan assays (primer/probe sets) (Applied Biosystems, Invitrogen) permitting the best coverage of the targeted gene BST2 Hs01561315_m1 was used for gene expression and further downstream in the factor analysis as it has the best coverage for the targeted gene.

A preamp pool of primers was prepared from equal volumes of each 20x TaqMan gene expression assays from the same gene expression assays to be used for qPCR and diluted by DNA suspension buffer (TE Buffer; 10 mM Tris, pH 8.0, 0.1 mM EDTA), Therefore each assay is at a final concentration of 0.2x (180 nM). Pre-amplification process was performed using limited fourteen cycles. cDNA prepared in a pre-amplification step allows for multiplex amplification of target genes were diluted 1:5 using TE Buffer.

Diluted cDNA was then loaded on a Fluidigm^®^ 96.96 Dynamic Array™ integrated fluidic circuit IFCs chip after priming using Fluidigm^®^ IFC controller HX respectively. TaqMan Gene Expression Assays were performed using the BioMark™ HD System and appropriate cycling protocols were applied for the 96.96 chip. The gene expression of cell markers and ISGs was analyzed by relative quantitative reverse transcription–polymerase chain reaction (qRT–PCR) using TaqMan reagents from Applied Biosystems. Gene expression data were normalized to peptidylprolyl isomerase A (cyclophilin A) gene (*PPIA*) as housekeeping reference gene and calculated using power to -ΔCt method. ISGs genes selected and IFN score was performed as previously described (El-Sherbiny et al. manuscript in review). We applied *PP1A* as house keeping gene

### IFN stimulation experiments

Peripheral blood was obtained from healthy donors, PBMCs prepared as described above or B-cells purified and then cultured in vitro as described before [6]. then cells were exposed to media alone or containing increasing levels of IFN-α, IFN-β, IFN-γ or IFN-λ for 48 hours before analysis of gene expression profile as described in gene expression studies and surface tetherin/CD317 protein levels on each subset of cells based on gating strategy.

### Statistical analysis

To investigate whether expression of the *BST2* gene was associated with surface protein levels measured through flow cytometry, values from SLE patients and healthy controls were pooled to give a wide range of values over which to compare the subsets. Gene expression values in ∆Ct were reflected so that the direction of effect was consistent for both gene expression and cell surface protein level whilst retaining log scaling; protein levels were ln-transformed prior to analysis. Pearson’s product-moment correlations were performed within each sorted cell subtype. In addition, gene expression and cell surface protein levels were ranked within individuals by cell subtype, to see whether consistent within-person ranking was obtained for each assay. Ranks of gene expression values and cell surface protein levels were assessed for agreement using quadratic-weighted Kappa (Kw) with confidence intervals estimated from 1000 bootstrapped samples.

A multilevel model was used to compare cell surface protein levels between SLE patients and healthy controls across differenT-cell subtypes identified during flow cytometry of PMBCs. Age at sampling was included as a covariate; estimated between-group differences were calculated at the mean age.

Interferon score based on gene expression levels were compared againsT-cell-specific BST2 protein levels to see which of these variables were more strongly associated with disease status. Analysis of covariance models, controlling for age, were constructed each of the cell subtype-specific protein levels, comparing patients with SLE to those with RA and to healthy controls. Effect sizes (partial eta squared) were calculated for each model to indicate which of the variables differed to the greatest extent between the different groups.

Within SLE patients, the variables were assessed for strength of correlation with total BILAG score and plasmablast count using Spearman rank correlation. To analyze association with disease activity the number of active BILAG domains (scoring A, B or C) was counted and was associated with interferon score and protein levels using multilevel modelling, controlling for age. Snijders/Bosker R-squared values (level 1)[53] were used to compare the strengths of association between the candidate assays. Within SLE patients, the variables were assessed for strength of correlation with plasmablast count, which was highly skewed in distribution, using Spearman rank correlation.

**Table. S1. Baseline Characteristics of SLE Patients.**

| Age (Mean (SD)) | 45.4 (14.3) | | | | |
| --- | --- | --- | --- | --- | --- |
| Female % | 91 | | | | |
| BILAG Organ Domain Scores (% patients) | **A** | **B** | **C** | **D** | **E** |
| Mucocutaneous | 5 | 18 | 16 | 44 | 18 |
| Musculoskeletal | 5 | 11 | 34 | 40 | 10 |
| Renal | 4 | 2 | 4 | 21 | 69 |
| Neuropsychiatric | <1 | 6 | 3 | 21 | 69 |
| Haematology | <1 | 4 | 46 | 37 | 12 |
| Cardiorespiratory | <1 | 2 | 2 | 24 | 72 |
| General | <1 | <1 | <1 | 37 | 61 |
| Gastro and Ophthalmic | <1 | <1 | <1 | 7 | 93 |

**Table S2. Differences between diagnosis groups in cell surface tetherin levels from cell subtypes**

| BST2/Tetherin protein level: | Geometric mean protein level* | | | SLE-RA | | SLE-HC | |
| --- | --- | --- | --- | --- | --- | --- | --- |
|  | SLE (n=53) | RA (n=12) | HC (n=16) | Ratio (90% CI), *P* value | Effect size* | Ratio (90% CI), *P* value | Effect size* |
| Monocytes | 3525 | 2572 | 2972 | 1.37 (1.00, 1.88), p=.103 | 0.03 | 1.19 (0.87, 1.61), p=.557 | 0.01 |
| T-cells | 718 | 398 | 506 | 1.80 (1.38, 2.37), p<.001 | 0.15 | 1.42 (1.09, 1.84), p=.029 | 0.06 |
| NK-cells | 1187 | 602 | 835 | 1.97 (1.50, 2.60), p<.001 | 0.18 | 1.42 (1.08, 1.85), p=.031 | 0.06 |
| Naïve B-cells | 1238 | 655 | 727 | 1.89 (1.41, 2.54), p<.001 | 0.14 | 1.70 (1.28, 2.26), p=.003 | 0.11 |
| Memory B-cells | 1722 | 898 | 1058 | 1.92 (1.46, 2.51), p<.001 | 0.17 | 1.63 (1.26, 2.11), p=.003 | 0.11 |
| Plasmablasts | 2860 | 1297 | 1799 | 2.20 (1.66, 2.93), p<.001 | 0.23 | 1.59 (1.21, 2.09), p=.006 | 0.09 |

*Partial eta squared

All values calculated at mean age.

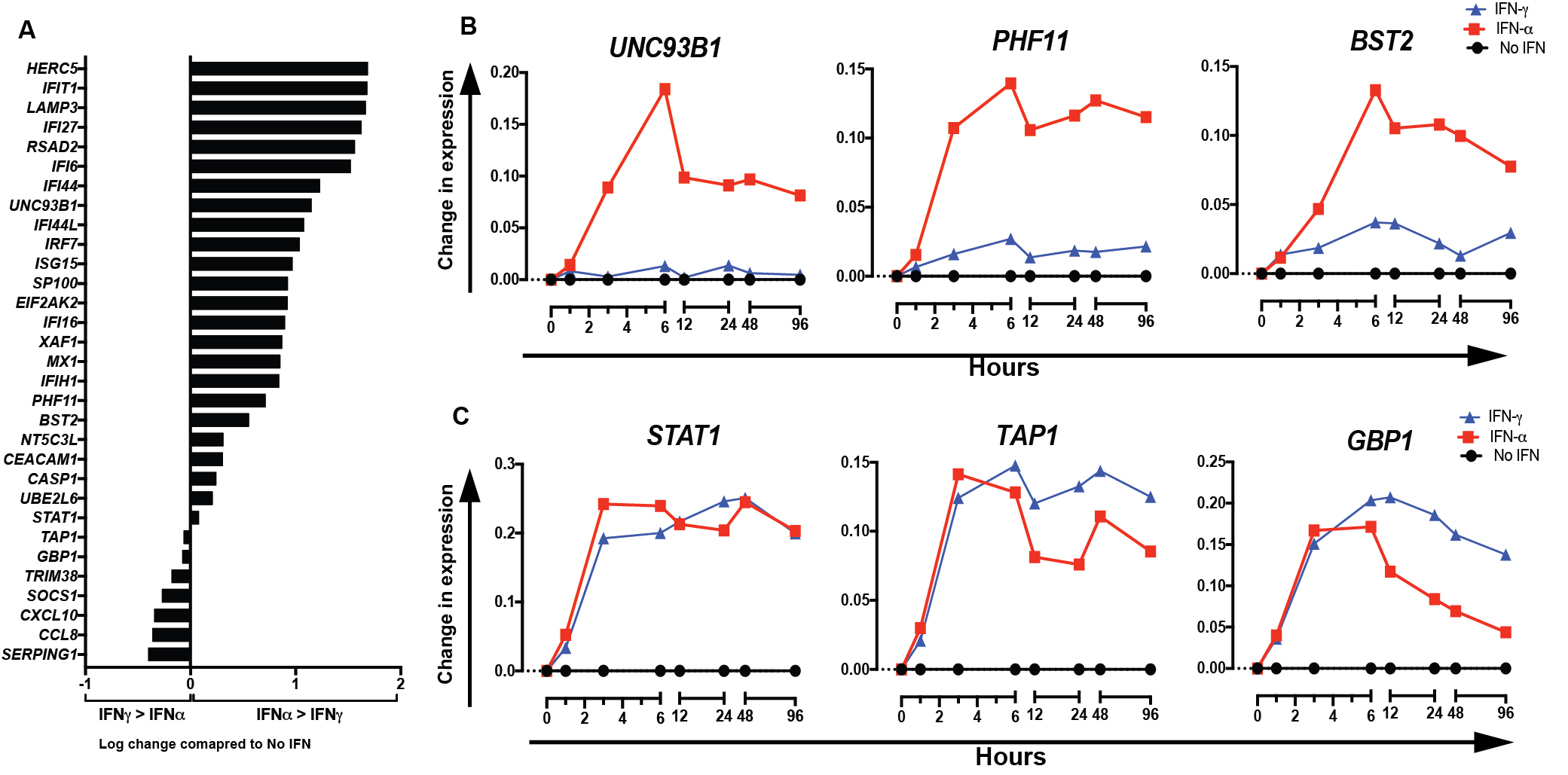

**Figure S1: Response of selected ISG expression after *in vitro* stimulation of B-cells**. Expression of selected ISGs was measured following in vitro stimulation of B-cells using either IFN-α or IFN-γ. B-cells were purified from peripheral blood and cultured in vitro as previously described (see methods). Activated B-cells were exposed to media alone or IFN-α or IFN-γ for between 1 and 96 hours before analysis of gene expression profile. (A) Shows log of ratio of increase in expression 6 hours after α vs. γ, each compared to no stimulation. Results are mean of 3 healthy donors. Values greater than 0 indicate greater increase in expression with α than γ. Values below 0 indicate greater increase in expression with γ than α. (B) Shows change in expression of 3 ISGs that predominantly respond to IFN-α, which include our flow cytometric marker, *BST2*/tetherin. (C) Shows change in expression for 3 ISGs, which do not demonstrate selective response to IFN-α in B-cells.

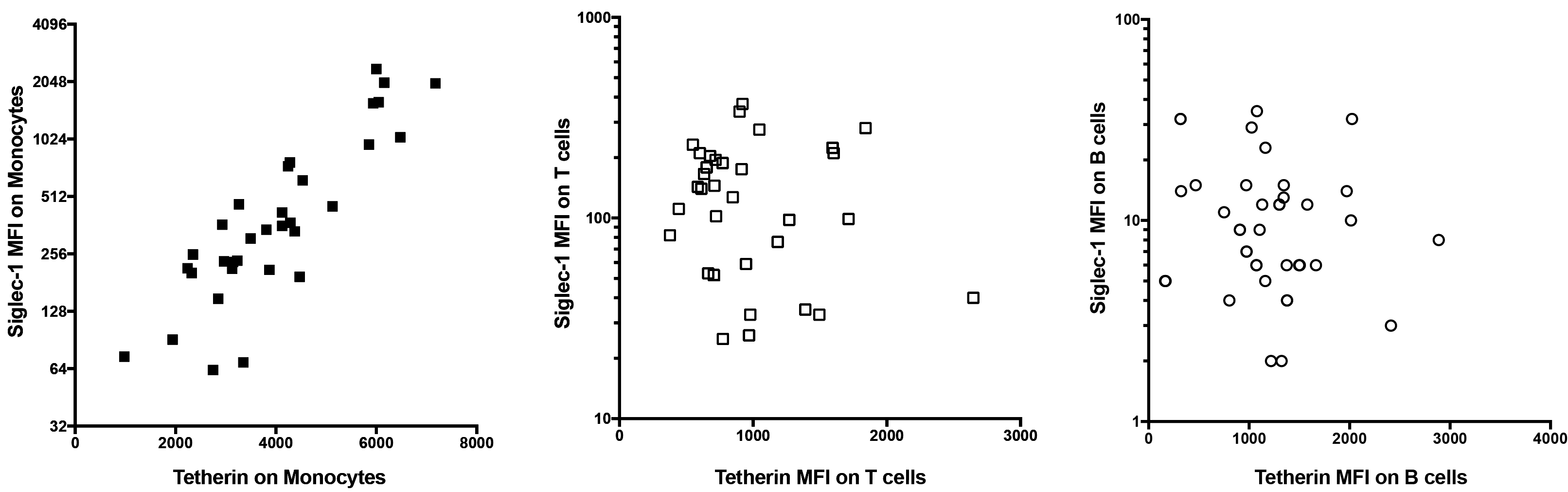

**Figure S2. Flow cytometry for tetherin (CD317) and Siglec-1 (CD169) mean fluorescence intensity on CD64+CD14+ monocytes, CD3+ T cells and CD19+ B cells.** Tetherin correlated closely with Siglec-1 on monocytes, in which both markers were expressed. As expected, Siglec-1 expression on T cells and B cells was minimal, and as such there was no correlation with tetherin.
